# Connectivity of variants in eQTL networks dictates reproducibility and functionality

**DOI:** 10.1101/515551

**Authors:** Sheila M. Gaynor, Maud Fagny, Xihong Lin, John Platig, John Quackenbush

## Abstract

Network analyses are a natural approach for identifying genetic variants and genes that work together to drive disease phenotypes. The relationship between SNPs and genes, captured in expression quantitative trait locus (eQTL) analysis, can be represented as a network with edges connecting SNPs and genes. Existing network methods treat such edges as fixed and known when they are most often thresholded estimates from eQTL regression. We consider various characterizations of an essential feature of nodes of eQTL networks, their degree centrality, that retains different data on eQTLs. We define the network metric of degree to be estimated by false discovery rates, test statistics, and p-values of the eQTL regressions in order to represent how central and potentially influential a SNP is to the network. We calculate degree metrics for data from 21 tissues from the GTEx project to assess the reproducibility, correlation across tissues, and, functional importance of degree.

## Introduction

Human diseases are influenced by many genetic variants that often act in concert to effect change in cellular function [10]. The overwhelming majority of these mutations lies in non-coding regions and are enriched within regulatory elements [1, 22, 7]. In particular, they are over-represented in one particular class of variant, expression quantitative trait loci (eQTL), that associate the presence of a genetic variant with the expression level of a gene [14, 5, 20, 15]. They have been demonstrated to play an important role in the causal pathway between genetic variants and disease [8], and further be shown to be enriched in certain tissues for particular diseases.

Principled methods are needed to explore the relationship between genetic variants, gene expression levels, and disease phenotype. Classical approaches such as genome-wide association studies (GWAS) and eQTL mapping studies identify relationships between variants and outcomes independently. These approaches consider only pairwise associations, and we are unable to use isolated association studies to elucidate the molecular mechanisms by which multiple genetic variants contribute to phenotype [22, 13]. In order to learn biological mechanisms of disease, integrative analyses of different types of genetic and genomic data is of increasingly significant importance and must improve to represent context-specific biological relationships and be reproducible across studies. We can identify genetic variants that influence cellular processes to alter phenotype by using network analysis to identify groups of genetic variants and genes that work together to collectively drive disease phenotypes.

Network analyses have emerged as an integrative approach to characterize complex genomic associations [3]. Bipartite networks are a natural representation for eQTL associations, where the edges between SNPs and gene expression represent the strength of the eQTL association [2, 18, 4]. Features of a network can inform function. For example, nodes that are more densely connected can represent natural divisions of functional relatedness. This representation has been shown to identify biological effects in chronic obstructive pulmonary disease (COPD) [9]. In COPD, GWAS-identified single nucleotide polymorphisms (SNPs) were found to be most central among groups of functionally related features [18, 6, 15].

Existing eQTL network approaches treat edges (such as the strength of eQTL associations) as known indicators, when they are in fact thresholded estimates from the initial eQTL analysis [1]. In this initial analysis, gene expression is regressed on SNP genotype to evaluate an eQTL association. After these associations are estimated, they are thresholded to obtain a binary edge, discarding potentially valuable data and introducing an additional source of error. This approach requires a relatively small computational burden as one may limit output to those meeting a minimum threshold and storing an incomplete network matrix without weights. One may posit that reducing the estimated regressions to dichotimized estimates to build a network may be detrimental to ensuring results are true and can detract from potential reproducibility. Methods that are more robust than operating on a threshold to account for the SNP-gene association from the eQTL network model can proposed to overcome these potential limitations. However, approaches that include fully weighted network representations have much greater computational burden given the need to retain and operate on output from millions of regression models. In this paper, we consider a set of edge representations of the SNP-gene association that can inform biological relationships, specifically towards estimating degree, a measure of how central a node is to the network. The degree is a measure of centrality that is associated with how essential a node is to function. We estimate degree metrics for eQTLs calculated from the Genotype-Tissue Expression (GTEx) project, allowing for the comparison of the eQTL findings across multiple bodily tissues. We characterize features of our defined degrees, consider their relationship to various functional features of the SNPs, and assess their reproduciblity.

## Methods

In this section, we describe our approach for constructing eQTL networks and defining the network metric of degree. This approach requires processed genotype and gene expression data, which can then be used to map eQTLs and build a network. We identify differences in the various approaches with regards to stability and computational feasibility. We also provide details of the implementation of these approaches and their reproducibility. An overview of the workflow is given in Figure 1.

**Figure 1:**
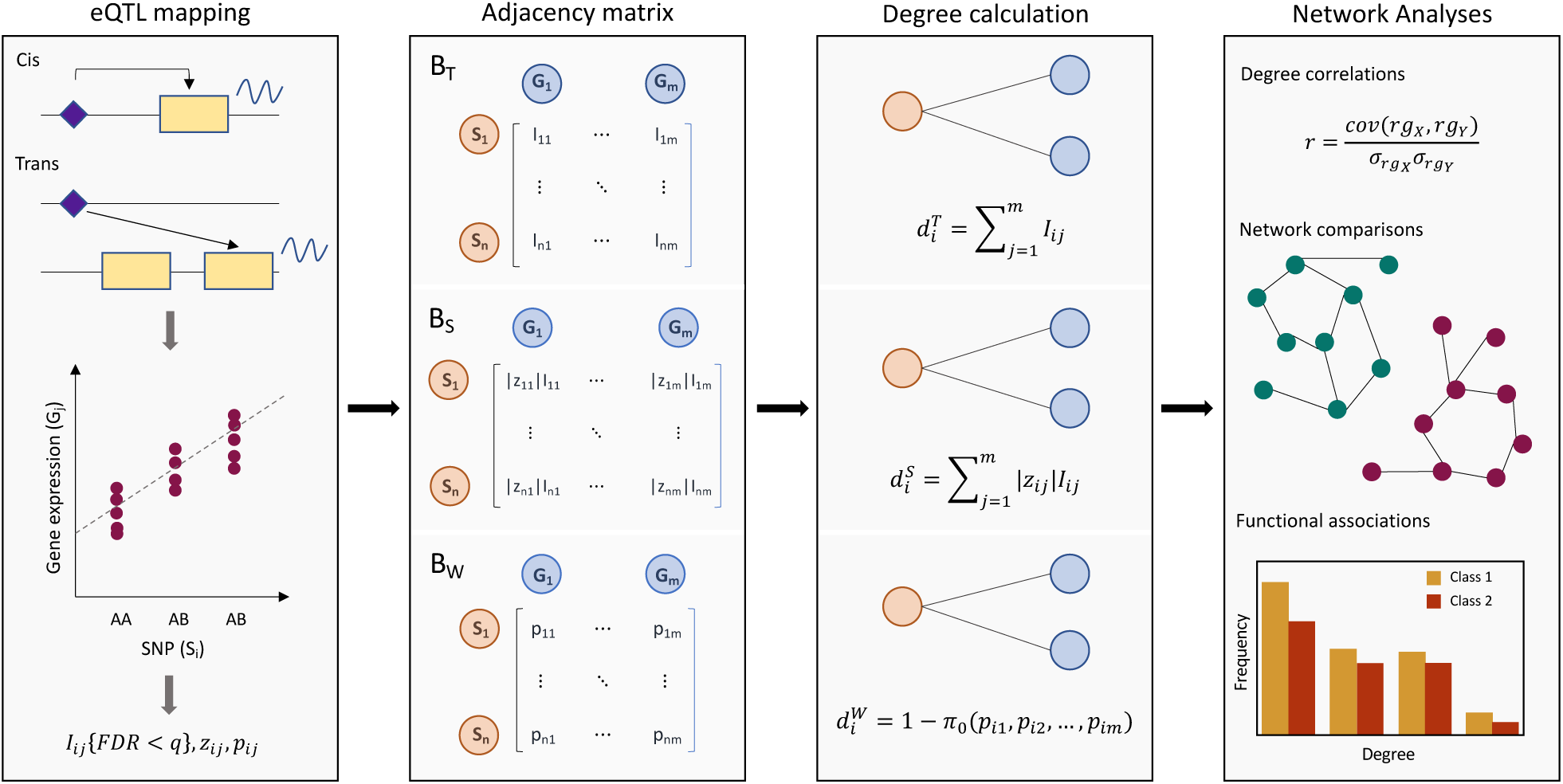
eQTL network construction and analysis workflow. eQTLs are mapped from genetic and gene expression data and a function of their associations is used to construct an adjacency matrix, from which network metrics such as degree can be calculated and used to infer scientific conclusions.

### 0.1 Network and degree definition

eQTLs are identified from the association between SNP genotypes and gene expression [12, 21]. Given *r* study observations, we have a matrix **S** of SNP genotypes and matrix **G** of gene expression, each with *r* rows representing observations and *n* and *m* columns respectively representing *n* SNPs and *m* genes. Consider some set of covariates **X**, such as principal components for population structure, sex, and age. For the eQTL regression of a particular SNP *i* on a locus’s gene expression *j* we have,

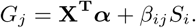

The collection of these eQTL associations can be represented as a bipartite network by casting a function of the association *β*_*ij*_ as an edge between SNP *i* and gene *j*. Specifically, we define an *N × N* adjacency matrix **A** to represent these connections,

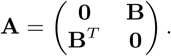

where **B** is a *n×m* matrix with rows representing SNPs, columns representing genes, and contains entries derived from the association of the SNP-gene pairings. The entries of **A** that would represent edges between the same node type (i.e. SNP-SNP relationships) are set to zero, as we are only considering the SNP-gene relationship. Previous studies on eQTL networks, such as Platig et al, defined the entries of **B** by dichotomizing all SNP-gene associations according to a cutoff *q* on the false discovery rate (FDR) for the eQTL regression, *I*_*i,j*_{*FDR <q*}. Thus when the estimated FDR of the eQTL regression was below the threshold of 0.2 for SNP node *i* and gene expression node *j* then *B*_*i,j*_ = 1, indicating there was an edge connecting the nodes, and *B*_*i,j*_ = 0 otherwise. We define this adjacency matrix representation to be **A**_*threshold*_. We consider two FDR estimation approaches. The first of which is genome-wide FDR calculation using the Benjamini-Hochberg procedure, considering all potential eQTLs. Secondly, we use the Benjamini-Hochberg procedure performed separately for *cis*-eQTLs and *trans*-eQTLs meeting a significance threshold. The Benjamini-Hochberg approach is considered appropriate as it allows for non-negative correlation between the tests.

We consider two alternative representations of the edges that retain more data and limit the influence of thresholding. Accordingly, we propose an adjacency matrix using a different representation of the SNPs and gene expression eQTL associations. First, we define an adjacency matrix that maintains the sparsity of **A**_*threshold*_ while incorporating the effect size. Calling this bipartite adjacency matrix **A**_*sparse*_, we define the entries of **B**_*sparse*_ to be *|z*_*i,j*_ *|I*_*i,j*_ {*FDR < q*}where *z*_*i,j*_ is the z-statistic for testing the eQTL regression parameter *β*_*i,j*_. Therefore when the estimated FDR of the eQTL regression was below the threshold of *q* for the SNP-gene pairing then *B*_*i,j*_ =*|z*_*i,j*_ *|* and *B*_*i,j*_ = 0 otherwise, providing a sparse representation incorporating the magnitude of the effect. We last define an adjacency matrix that does not threshold any aspect of the association, **A**_*weight*_ Here, we define the entries of **B**_*weight*_ to be *p*_*i,j*_ where *p*_*i,j*_ is the p-value for the z-test of the eQTL regression parameter *β*_*i,j*_ between SNP *i* and gene *j*. This is a dense representation that includes the p-value of all eQTL associations. These three adjacency matrix representations are shown below for the **B** partition of the adjacency matrix **A**. We consider three values for *q* in our applications, 0.05, 0.10, and 0.20.

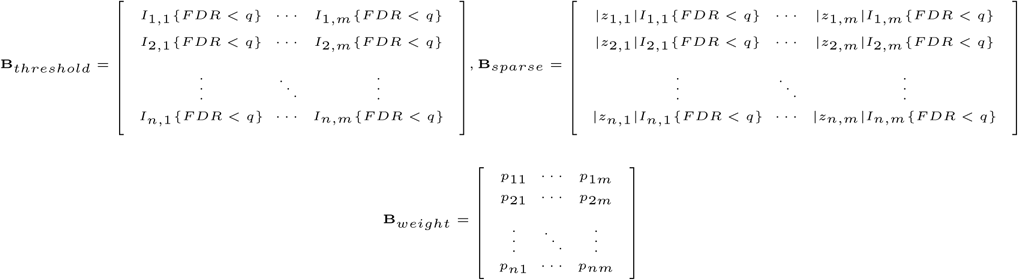

We are particularly interested in identifying SNPs (or genes) that are central to the network by comparing the network metric of degree, which is a measure of node centrality. For an eQTL network, a SNP with high degree is most highly connected to the expression of genes and therefore should be highly functionally relevant. We consider representations of the SNP-level network metric degree particular to each of the defined adjacency matrix. In particular, for SNP 1 we define

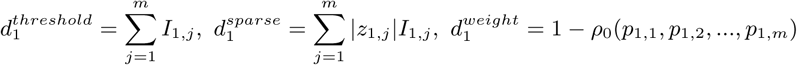

For the thresholded adjacency matrix **A**_*threshold*_, we take the standard row summation to obtain a count of the number of (binary) connections a SNP has to genes, which we abbreviate as *d*^*t*^. The sparse adjacency matrix **A**_*sparse*_ has a weighted version of this sum, incorporating the magnitude of the test statistic, and is referred to throughout as *d*^*s*^. For **A**_*weight*_, we estimate the proportion of significant eQTL analyses for a particular SNP, or the proportion of genes whose expression are influenced by the SNP by utilizing the proportion of true null hypotheses, *ρ*0. Thus SNPs that have higher degree if they are estimated to have fewer true null associations to the genes. This degree 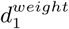, or simply *d*^*w*^, thus equires the estimation of the proportion of null hypotheses. The proportion is estimated utilizing an empirical method of estimating the proportion of true null hypotheses. The method of Heller and Yekutiel is used, where the marginal density of the test statistics is modeled via Poisson regression. The proportion of nulls is estimated by assuming that the statistics falling in the central 50% of the null distribution are null. This has been implemented in the R package `locfdr`.

### 0.2 Application to GTEx Study

Data from the NHGRI Genotype-Tissue Expression (GTEx) project was used to build eQTL networks across tissues. The GTEx project is a consortium collecting genotype and expression data from multiple human tissues from hundreds of human donors. We downloaded the Version 7.0 whole genome sequencing and RNA-seq data from the database of Genotypes and Phenotypes (dbGaP) under accession phs000424.v7.p2. A threshold of at least 200 individuals per tissue available was considered for appropriate statistical power and network stability; sex-specific tissues were n ot included. Computations on the GTEx data were r un on t he B ridges system at the Pittsburgh Supercomputing Center (PSC) and the Odyssey cluster supported by the FAS Division of Science, Research Computing Group at Harvard University. The sequencing data were processed in Plink 1.90 to only retain SNPs, and remove variants with genotype missingness greater than 10% or minor allele frequency less than 0.05. The RNA-Seq data were processed using the `YARN` package [16] in Bioconductor and normalized using `qsmooth` [11] in Bioconductor; extraction effects were adjusted for using the `limma R` package [19]. Genes with a read count of less than 5 were not considered to be expression; genes expressed in less than 10 samples or considered a pseudogene by the Ensembl database in `biomaRt` were excluded.

### 0.3 eQTL Mapping

We used a linear regression model with covariates assuming an additive effect of genotypes to map eQTLs. We accounted for population stratification by using the first three principal components of the genotypes as covariates. We further adjusted for sex, age, and genotyping platform, with the model for gene *j* and SNP *i* given by,

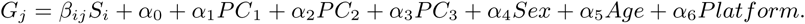

The regression coefficient *β*_*ij*_ was then used in network estimation as previously described. Wald tests were used for performing inference on *β*_*ij*_. Genes were mapped using the biomaRt package in Bioconductor. We defined *cis*-eQTLs to be SNPs associated with genes where the distance between them is less than or equal to 1Mb pairs of each other; all other SNP-gene pairings were defined as *trans*-eQTLs. Analyses were conducted in R 3.3.0 and utilized the `MatrixEQTL` and `locfdr` packages. All calculations were massively parallelized across SNPs. The eQTL mapping by the GTEx Consortium was compared by downloading the single-tissue *cis*-eQTL results for significant variant-gene associations based on permutations from gtexportal.org.

### 0.4 Correlation

We consider the correlation of the SNP degree in two settings: across different tissues and within a particular tissue. We can compare the degree of SNPs between tissues via correlation to define the network-level relationship between tissues. We expect that, particularly for *cis*-eQTLs, that tissue-specific networks should share features. Given the non-normal distributions of each of the degree measures, we use Spearman correlation. We further consider the correlation of the SNP degree within a particular tissue, predominantly as a demonstration of reproducibility for each degree measure. We perform sample splitting in the smallest and largest tissue tissue by randomly splitting the observations in half, constructing networks and calculating the degree, and then estimating the correlation of the degree between the splits. This was repeated five times i n e ach t issue to account for variability.

### 0.5 Gene Networks

We built tissue-specific gene co-expression networks to consider relationships between genes. We use the Weighted Gene Co-expression Network Analysis as implemented in the `WGCNA R` package. This approach requires the selection of a soft thresholding power for constructing the network, which was selected based on inspecting by plot the first inflection point for the scale-free topology fit index curve. The co-expression network was constructed using all of the genes considered in eQTL mapping.

We also built tissue-specific regulatory networks i n the same approach as Sonawane et al. In particular, we used the Passing Attributes between Networks for Data Assimilation (PANDA) approach to build a gene regulatory network using the `pandaR` package in Bioconductor. We used the motif and protein-protein interaction data provided in Sonawane 2017 which were derived from the Catalog of Inferred Sequence Binding Preferences and StringDb; further details are given in the Experimental Procedures. We used the present gene expression data in the PANDA approach to identify regulatory networks that relate transcription factors and genes. We calculated the degree of genes in the regulatory network using the transformation to the edge weights suggested in Sonawane to account for negative edge weights, specifically for a given network that

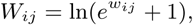

where *w*_*ij*_ is the edge weight between transcription factor *i* and gene *j*.

We compare these networks to the eQTL networks by considering the degree of genes, rather than SNPs. We use the same approaches to degree estimation in the networks, but rather than considering the number of genes connected to a particular SNP we fix the gene and consider the relationship of all SNPs to the gene. We can thus compare the degree across the different types of gene networks, towards understanding the differences in the relationships that they capture.

### 0.6 Functional modeling

We downloaded functional annotations from the Combined Annotation Dependent Depletion (CADD) database. We mapped our variants to CADD v1.2 given that the GTEx data is based on the GRCh37/hg19 genome build; all variants were available in CADD. This included a variety of different annotation features; we considered only those with limited missingness.

In order to model the association between functional annotations and tissue-specific network degree, we used a Poisson generalized estimating equation (GEE) model. We consider the SNP-level measure of degree using the thresholded definition *d*^*threshold*^, excluding any isolated nodes or SNPs with degree equal to zero. We shift the degree measures down by one to have the interpretation of additional connections. The SNPs have not been pruned for linkage disequilibrium; to account for this correlation we calculated haplotype blocks in Plink 1.90 with a maximum block size of 5-Mb. Blocks were then treated as clusters in the GEE model using an exchangeable working correlation. The model is thus given for a specific tissue as,

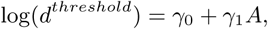

where *A* is a particular annotation value, either continuous or binary.

## Results

### eQTL networks are dependent on edge definition

We considered eQTL networks constructed from the genotype and RNA-Seq data for 21 tissue types with adequate sample sizes from the GTEx Version 7.0 dataset, as described in the Methods. After data processing primarily to limit the analysis to common variants and normalize the expression data, we retained 5,442,245 SNPs for all observations and 25,305 genes on average across tissues. The tissues included in our analysis had between 203 and 491 individuals.

Exhaustive eQTL mapping adjusting for sex, age, genotyping platform and the first three principal components was performed for each of the 21 prepared tissues. *cis*-eQTL mapping was performed for variants within 1 Mb of a gene. We identified a minimum of 221786 (stomach) and maximum of 810853 (thyroid) *cis*-eQTLs at a false discovery rate (FDR) of less than 0.05. *trans*-eQTL mapping was performed separately for variant-gene pairings outside the 1 Mb window, for which we identified between 57621 (stomach) and 142369 (thyroid) SNP-variant pairings at an FDR threshold of 0.05. We validated our eQTL mapping results to those reported by the GTEx Consortium, which reported *cis*-eQTL associations. We found that on average 78% (sd=2%) of our *cis*-eQTL calls were also in the GTEx Consortium results. Whole blood had the lowest percentage of *cis*-eQTLs also identified by the GTEx Consortium at 73%; as many 80% of *c is*-eQTLs were also identified in our results as in artery tibial. The eQTL mapping results are summarized in Table A1 of the Appendix.

We constructed eQTL networks from the eQTL association results based on a variety of edge definitions. The definitions varied primarily as to whether they considered SNP-gene pairings on a genome-wide or location-specific scale and whether the edges were weighted. Given the edge definitions, we calculated the proposed degree metric for each SNP. We then accounted for gene location by calculating the degree metrics for *cis*-eQTLs and *trans*-eQTLs separately for the approaches allowing for location specificity. The network construction was performed in a massively parallelized approach for each of the 21 tissues.

The distributions of the different degree metrics across tissues is given in Figure 2. Panels A-C demonstrate the non-zero degree distribution for the *d*^*t*^ genome-wide, *d*^*t*^ location-specific, and *d*^*w*^ genome-wide approaches. As expected given the limited amount of eQTLs across the genome, most SNPs had degree equal to zero in each of the approaches. For the genome-wide thresholded *d*^*t*^(and thereby sparsely weighted *d*^*s*^) approaches, 88-92% of SNPs on average across each of the tissues had degree equal to zero; 91-93% of SNPs on average across each of the tissues in the location-specific approaches had degree equal to zero. The genome-wide *d*^*w*^ approach had 72% non-zero degree SNPs for a tissue on average. The weighted degree has a lower percentage of SNPs with degree equal to zero, ranging from 69-77% across tissues. The thresholded degree estimates are highly correlated with the sparsely weighed degree (*ρ >* 0.99), as suspected given that they rely on the same indicator. It is excluded from Panel E, as *d*^*t*^ and *d*^*s*^ rely on the same indicator and thus have the same proportion of zero degree SNPs. As demonstrated in Figure 2, the distributions are highly skewed to the right while excluding the zero degree SNPs.

**Figure 2:**
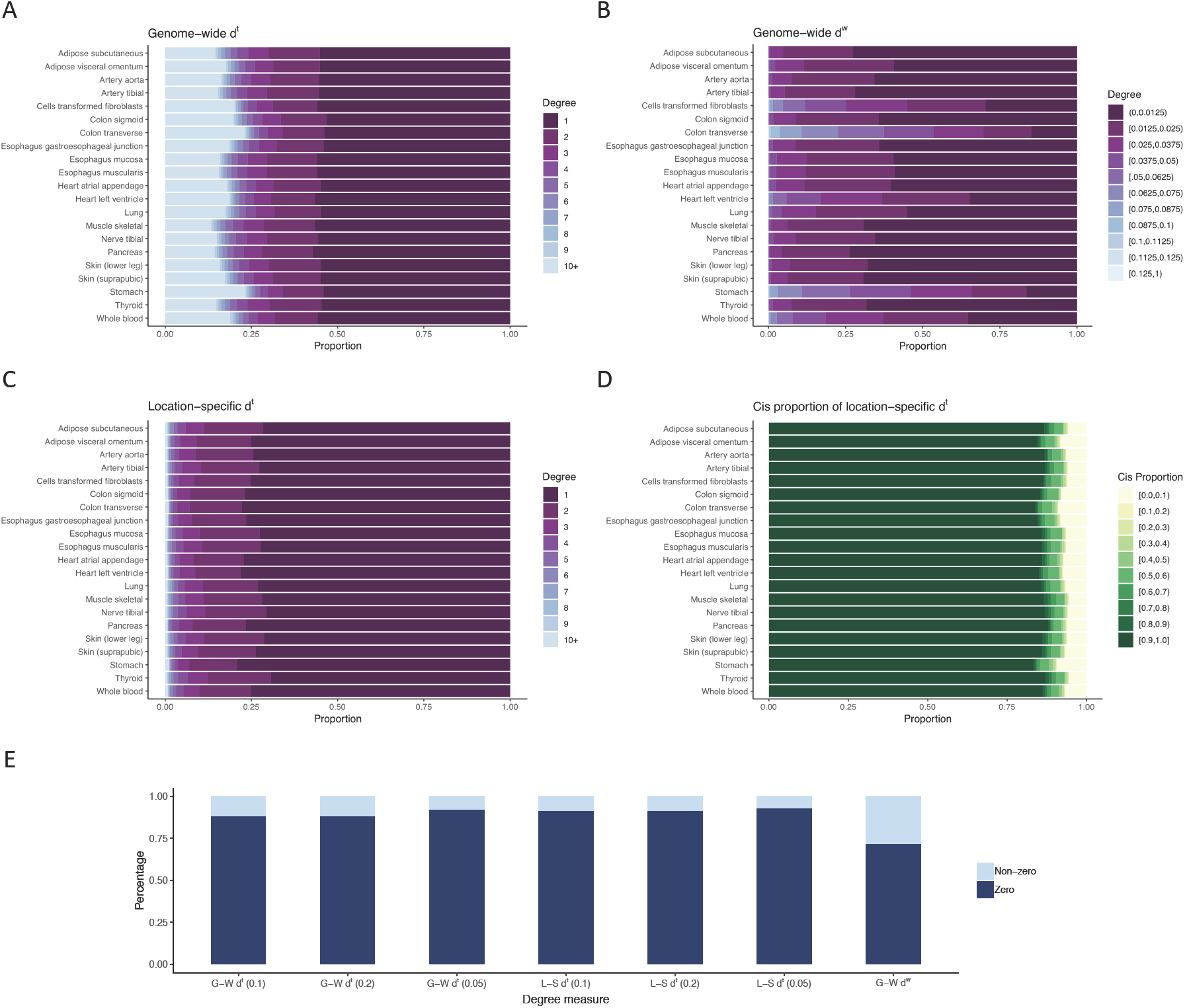
Distributions of degree measures. The distribution of non-zero degrees is given in panels A, B, and C using the thresholded approach genome-wide, weighted approach genome-wide, and thresholded approach location-specifically. The proportion of non-zero degree measures that are attributable to *cis*-eQTL associations in the location-specific thresholded approach is given in panel D. Panel E shows the proportion of non-zero degrees under the location-specific and genome-wide approaches with various thresholds.

Consistent with previous findings that *cis*-eQTLs are more commonly identified than *trans*-eQTLs, we observe that the majority of non-zero SNP-gene edges are *cis*-eQTLs, as shown in Panel D. For the majority of SNPs with non-zero degree, the *cis* component constitutes 100% of the non-zero degree magnitude. Variants with regulatory functions tend to act locally within the megabase window, leading to higher proportions of *cis*-eQTLs than *trans*-eQTLs. The proportions are consistent between the thresholded and sparsely weighted degree metrics though shown only for the location-specific thresholded approach.

Figure 2 also illustrates the greatly increased computational burden of the different estimation approaches. For the thresholded and sparsely weighted degree, only SNP-gene associations that meet the particular threshold must be stored intermediately and degree is calculated via simple summations by SNP. Given the high proportion of zero degree SNPs, this means oftentimes the association will not need to be stored in memory and one can capitalize upon current eQTL software that allows for efficient univariate regression for large datasets. Alternatively, for the weighted degree we must calculate and retain in computer memory all SNP-gene associations in order to perform more computationally intensive SNP-level estimation. The resulting output from each of the approaches is of equal computational cost; however, in order to obtain the degrees in a fully weighted approach one must retain and sort through all associations.

We compared the computational impact of calculating the degree of 10,000 SNPs for our considered set of 24,634 genes in the largest tissue (skeletal muscle). The run time was based on a 2.70 GHz laptop with 16Gb of memory. We observe that the location-specific degree computation is more than five times faster than the genome-wide computation. This is significant considering that studies are currently growing in the number of SNPs genotyped as well as those that can be imputed with high confidence, therefore scalability is of high importance.

### Degree definition determines stability

#### Within-tissue reproducibility

The stability of eQTL networks and their metrics is dependent on selecting appropriate network definitions and having an appropriate sample size. The stability of estimating SNP degree in independent samples and the impact of reduced sample sizes were evaluating by splitting the tissues, computing the degree metrics, and assessing their concordance between the split sample estimates. For the smallest and largest tissues, colon sigmoid (n=203) and skeletal muscle (n = 491), we randomly split the data in half. We estimated the degree measures on each half of the split data, and calculated the Spearman correlation between them. This was repeated and averaged across 5 subsamples to account for variability. The estimated correlations for the thresholded, sparsely weighted and weighted degree under both the genome-wide and location-specific settings are given in Table 1 for all *q* threshold values.

**Table 1:** Correlation of estimated SNP degree under all degree definitions between sample splits of colon and skeletal muscle tissues. All correlations are averaged across five sample splits.

|  | Degree Measure | Colon - Mean (SD) | Skeletal Muscle - Mean (SD) |
| --- | --- | --- | --- |
| Genome-wide | $d^t, \sum_i \mathbb{1}_{\{FDR < 0.05\}}^i$ | 0.121 (0.003) | 0.235 (0.003) |
| | $d^t, \sum_i \mathbb{1}_{\{FDR < 0.1\}}^i$ | 0.078 (0.002) | 0.154 (0.002) |
| | $d^t, \sum_i \mathbb{1}_{\{FDR < 0.2\}}^i$ | 0.050 (0.002) | 0.092 (0.001) |
| | $d^w, 1 - \pi_0$ | -0.001 (0.002) | 0.0003 (0.002) |
| | $d^s, \sum_i Z_i \cdot \mathbb{1}_{\{FDR < 0.05\}}^i$ | 0.126 (0.003) | 0.246 (0.003) |
| | $d^s, \sum_i Z_i \cdot \mathbb{1}_{\{FDR < 0.1\}}^i$ | 0.083 (0.001) | 0.165 (0.002) |
| | $d^s, \sum_i Z_i \cdot \mathbb{1}_{\{FDR < 0.2\}}^i$ | 0.053 (0.002) | 0.102 (0.001) |
| Location-specific | $d^t, \sum_i \mathbb{1}_{\{FDR < 0.05\}}^i$ | 0.541 (0.008) | 0.603 (0.005) |
| | $d^t, \sum_i \mathbb{1}_{\{FDR < 0.1\}}^i$ | 0.490 (0.009) | 0.549 (0.005) |
| | $d^t, \sum_i \mathbb{1}_{\{FDR < 0.2\}}^i$ | 0.407 (0.006) | 0.469 (0.003) |
| | $d^s, \sum_i Z_i \cdot \mathbb{1}_{\{FDR < 0.05\}}^i$ | 0.544 (0.008) | 0.611 (0.005) |
| | $d^s, \sum_i Z_i \cdot \mathbb{1}_{\{FDR < 0.1\}}^i$ | 0.494 (0.009) | 0.559 (0.005) |
| | $d^s, \sum_i Z_i \cdot \mathbb{1}_{\{FDR < 0.2\}}^i$ | 0.413 (0.006) | 0.481 (0.003) |

We observe on average a small correlation between sample splits in both tissues in the genome-wide measures, with the correlations ranging from 0.05 to 0.24 for the *d*^*t*^ measures. The correlation increases slightly when incorporating the measure of effect, *Z*_*i*_, in the sparse degree *d*^*s*^, where the correlation ranges from 0.05 to for the thresholded measure. The average correlation between the splits decreases with increasing FDR threshold *q*, which is also observed in the location-specific measures. The weighted degree has notably lower correlation in the subsamples, in fact slightly negative correlation for colon sigmoid, indicating that it is a less consistent measure of degree than the other two degree measures considered. The location-specific measures have notably higher correlation on average between sample splits than the genome-wide approaches; the average correlation between splits for *d*^*t*^ ranges from 0.41 to 0.6. Again, the correlation increases very slightly when sparsely weighting.

Previous eQTL studies and power calculations have demonstrated that typically larger sample sizes than these subsamples are required for confidently mapping eQTLs. This moderate concordance between the subsample and full sample degree metrics illustrate the lack of stability of an eQTL network in a small network. The correlation between the sample splits increases on average between colon and skeletal muscle, which is likely attributable to having more than double the sample size for skeletal muscle.

#### Cross-tissue correlation

We compared the degrees identified in the tissue-specific networks using Spearman correlation; we expect moderate correlation across all tissues, particularly given that *cis*-eQTLs contribute a high proportion of the non-zero degree measures and are more often replicated across tissues. Figure 3 provides a summary of the trends observed in this correlation analysis. We observe higher correlation for the location-specific degree measures than the genome-wide measures; we note that the multiple testing adjustment allows for the less tissue-specific *cis*-eQTLs to be more readily identified than in the genome-wide setting. The mean Spearman correlation for the SNP degree is 0.20 for all tissue pairs for the genome-wide *d*^*t*^ with threshold 0.1; the mean is 0.47 for the location-specific *d*^*t*^ with threshold 0.1. In the weighted degree setting *d*^*w*^, the average correlation for pairs of tissues was 0.016, with a range of −0.004 to 0.056. This further suggests that this method, based on the estimation of the proportion of null hypotheses, is not reliable as we expect positive correlation between tissues.

**Figure 3:**
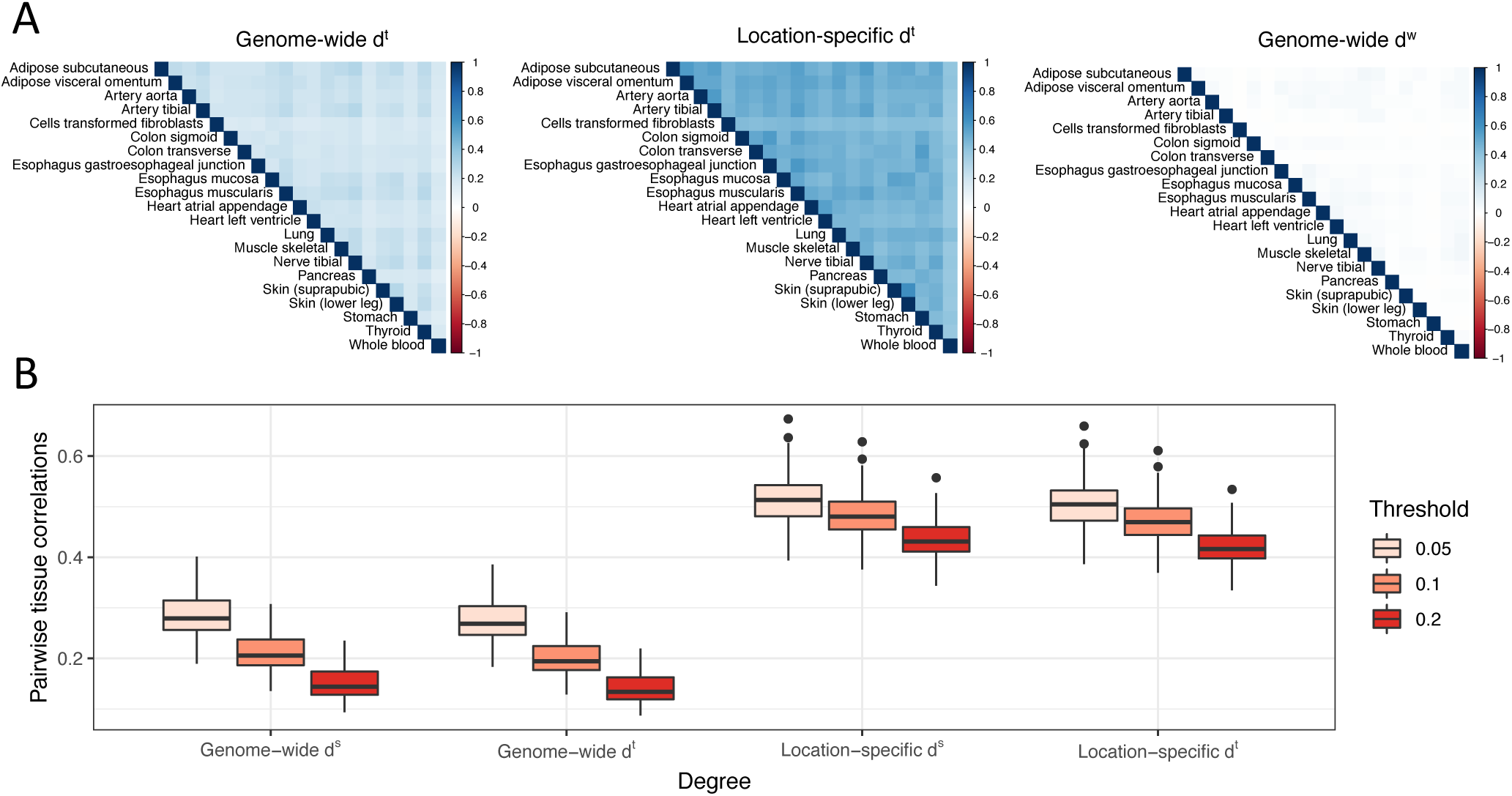
Degree correlations between tissues. The pairwise correlations between tissues under three degree definitions, thresholded with *q* = 0.1 and weighted, are given in Panel A. Panel B shows the distribution of the pairwise correlations considering different *q* thresholds across degree definitions.

For all levels of thresholding and for both genome-wide and location-specific definitions, the correlation between tissues is slightly increased from the thresholded degree *d*^*t*^ to the sparse degree *d*^*s*^. This was also observed in the within-tissue setting. Holding the threshold value and location consideration constant, the difference in average correlation across tissues between *d*^*s*^ and *d*^*t*^ ranges from 0.009 to 0.015. Higher, or more relaxed, FDR thresholds led to lower average pairwise correlations between tissues as evidenced by Panel B of Figure 3. A potential contributor may be that given the relaxed threshold, there is an increased potential for identifying sub-threshold tissue-specific *trans*-eQTLs as *trans*-eQTLs have lower power for eQTL mapping than *cis*-eQTLs.

### Degree correlation with other gene networks

We further calculated the degree of genes in the eQTL network, following the same degree definitions but estimating the relationship of all SNPs to a single gene, rather than all genes to a single SNP. We first assessed the correlation of the gene degree in the eQTL network to the in-degree, out-degree, and total degree of the WGCNA network. Considering the Spearman correlation between these degree measures, all associations were significant when considering the location-specific thresholded degree with the threshold *q* = 0.10. Most tissues maintained significant degree correlations when only considering *cis*-eQTL networks. Alternatively, the majority of tissues did not have significantly correlated degree measures between the eQTL network degree and the PANDA networks. These incongruent findings are consistent with the notion that the co-expression and regulatory networks capture different biological features.

### Degree correlates with different functional characterizations

We annotated the SNPs in each tissue with the Combined Annotation Dependent Depletion (CADD) score in order to explain SNP functionality. We considered a small set of annotations selected based upon missingness in the annotations and relatedness to eQTLs; we modeled their association with the location-specific thresholded degree *d*^*t*^ with the threshold *q* = 0.10. Table 2 provides the proportion of associations between the annotation measure and SNP degree that were found to be significant in the 21 tissues considered, with complete annotation description given in Table A2 in the Appendix. We consider the SNP degree in the context of the complete eQTL network and the eQTL network constructed only from *cis*-eQTLs.

**Table 2:** Proportion of tissues with *d*^*t*^ (*q* = 0.1) associated with a particular functional annotation. The proportion is given for both the complete eQTL network and the *cis*-eQTL only network.

|  | Proportion significant (all eQTL) | Proportion significant (*cis*-eQTL) |
| --- | --- | --- |
| CpG | 50.00 | 40.00 |
| priPhCons | 10.00 | 5.00 |
| mamPhCons | 0.00 | 0.00 |
| verPhCons | 0.00 | 0.00 |
| GerpN | 15.00 | 45.00 |
| GerpS | 0.00 | 0.00 |
| bStatistic | 100.00 | 90.00 |
| mutIndex | 75.00 | 45.00 |
| dnaHelT | 0.00 | 0.00 |
| dnaMGW | 0.00 | 0.00 |
| dnaProT | 45.00 | 15.00 |
| dnaRoll | 0.00 | 0.00 |
| EncH3K27Ac | 0.00 | 10.53 |
| EncH3K4Me1 | 15.79 | 0.00 |
| EncH3K4Me3 | 5.26 | 5.26 |
| EncNucleo | 0.00 | 15.79 |
| minDistTSS | 50.00 | 61.11 |
| minDistTSE | 66.67 | 41.67 |
| RawScore | 25.00 | 8.33 |
| PHRED | 25.00 | 8.33 |

We observe that some functional annotations are significantly associated with the SNP degree. The percent CpG within a 75bp window was associated with degree in half of the tissues; CpG sites that are differentially methylated have been suggested to mediate the relationship between genetic variation and expression [17]. We annotated the SNPs with the maximum ENCODE H3K4 methylation level, which was significantly positively associated with the degree in a number of tissues. This methylation level is a hallmark of primed enhancers, and eQTLs are known to be enriched in enhancer and promoter regions.

Given that not all annotations are associated with the eQTL SNP degree, this is an indication that it may represent a different aspect of the biological process. The significant associations do suggest that SNPs that may have a more significant and potentially detrimental impact through deleteriousness are not highly correlated with high degree, which may be because SNPs with potentially large detrimental effects may be less central in favor of SNPs with more consistent behavior.

## Discussion

We have proposed a set of approaches for defining eQTL networks and corresponding degree metrics. We applied our method to 21 tissues from the GTEx study. Each of the degree metrics has a different computational burden. The thresholded and sparsely weighted measures require notably less computational resources as they can be simply computed via summations from thresholded eQTL mapping results. The weighted degree measure requires one to exhaustively output all eQTL relationships and then perform further estimation on all output. All of these degree measures can be completely parallelized for optimal computation, but the impact is nonetheless an important consideration as one often would seek to perform these analyses genome-wide.

We find that all of the degree distributions are highly skewed, which is expected as mapped eQTLs are sparse across the genome. In our analysis of GTEx tissues, we find that degrees are correlated across tissues particularly for *cis*-eQTLs. Most of the contributions to the degree measure come from *cis*-eQTLs, as is expected because they are more commonly identified than *trans*-eQTLs. The *trans*-eQTLs are not significantly correlated, which is further evidence that *trans*-eQTLs are highly tissue specific. The correlation across tissues using the weighted degree method does not demonstrate any estimated correlation between tissues. One may expect this to be more likely than for the thresholded and sparsely weighted degree measures as the weighted degree is a more dense distribution. There is further explanation in the estimation approach of the proportion of null hypotheses; given the sparsity of signal amongst a large number of tests the measure does not perform well.

There was an overrepresentation of high degree SNPs in promoter region than those that are not in promoter region. We indeed expect SNPs that are in promoter regions and enhancer elements to have higher degree. We further demonstrate through other annotations that SNPs at higher risk of having more detrimental impact-such as being deleterious-are less connected. They are likely less central because in the event that the variant is indeed deleterious, it can have a significant and potentially harmful impact on the cell.

Our results demonstrated a general lack of consistency in results and concordance in degree specifically when using the weighted approach. This may be due to the fact that the other two methods allow for more false positive contributions to the degree calculation given the relaxed FDR threshold in calculation. Further, these methods separate between *cis*- and *trans*- eQTLs when calculating the FDR rates as it has been biologically evidenced that these two eQTLs have very different prevalences and allows for separate treatment. The weighted approach considers all eQTLs together and as a continuous measure of the could then be considered more stringent.

We have been able to characterize the degree of SNPs in eQTL networks under three different adjacency matrices. We observe more stable and expected results under the thresholded and sparsely weighted degree metrics, in addition to benefiting from efficient computation. The fully weighted approach, while statistically pleasing, does not capture the distinction between *cis*- and *trans*-eQTLs as the other two approaches and we observe less consistent performed. This may be further attributable to challenges in precisely estimating the proportion of null hypotheses. Further work would include pursuing fully weighted representations of the eQTL network while calculating an estimate of the proportion of null for a SNP stratified by *cis*- and *trans*-eQTLs. Additionally, it would be of interest to apply this framework to other biological QTL networks and further allow for comparisons across QTL networks.

## Acknowledgements

This work was supported by the National Science Foundation Graduate Research Fellowship (DGE1144152) and by the National Institutes of Health F31 (HL138832-01). This work was conducted under dbGaP approved protocol 9112.

## Appendix

**Table A1:** eQTL findings from the present analysis and the GTEx consortium results.

| Tissue | Samples | *cis*-eQTLs<br>(GTEx) | *cis*-eQTLs<br>(Present) | *cis*-eQTLs<br>(Shared) | Proportion | *trans*-eQTLs<br>(Present) |
| --- | --- | --- | --- | --- | --- | --- |
| Adipose subcutaneous | 385 | 1564953 | 638876 | 507250 | 0.79 | 115341 |
| Adipose visceral omentum | 313 | 1038696 | 411273 | 323017 | 0.79 | 89520 |
| Artery aorta | 267 | 1080810 | 463015 | 361323 | 0.78 | 88424 |
| Artery tibial | 388 | 1609924 | 585594 | 470809 | 0.80 | 103612 |
| Cells transformed fibroblasts | 300 | 1328237 | 392719 | 314136 | 0.80 | 82504 |
| Colon sigmoid | 203 | 697429 | 295953 | 224757 | 0.76 | 61283 |
| Colon transverse | 246 | 832982 | 258587 | 211486 | 0.82 | 63682 |
| Esophagus gastroesophageal junction | 213 | 733134 | 308341 | 238498 | 0.77 | 68730 |
| Esophagus mucosa | 358 | 1541480 | 581180 | 462245 | 0.80 | 109019 |
| Esophagus muscularis | 335 | 1447438 | 580662 | 460083 | 0.79 | 110732 |
| Heart atrial appendage | 264 | 878468 | 373623 | 282789 | 0.76 | 72055 |
| Heart left ventricle | 272 | 777906 | 317596 | 239656 | 0.75 | 66506 |
| Lung | 383 | 1464047 | 521242 | 418495 | 0.80 | 102589 |
| Muscle skeletal | 491 | 1496009 | 637971 | 499771 | 0.78 | 107236 |
| Nerve tibial | 361 | 1922195 | 740902 | 589621 | 0.80 | 126104 |
| Pancreas | 220 | 728858 | 387287 | 286149 | 0.74 | 70967 |
| Skin (not sun exposed suprapubic) | 335 | 1306368 | 500292 | 397191 | 0.79 | 95821 |
| Skin (sun exposed lower leg) | 414 | 1772181 | 679272 | 543205 | 0.80 | 123795 |
| Stomach | 237 | 636426 | 221786 | 176756 | 0.80 | 57621 |
| Thyroid | 399 | 2112918 | 810853 | 657095 | 0.81 | 142369 |
| Whole blood | 369 | 1052542 | 340837 | 270676 | 0.79 | 69559 |

**Table A2:** Complete description of features used from CADD database in functional analysis

|  | Description |
| --- | --- |
| CpG | Percent CpG in a window of +/- 75bp |
| priPhCons | Primate PhastCons conservation score (excl. human) |
| mamPhCons | Mammalian PhastCons conservation score (excl. human) |
| verPhCons | Vertebrate PhastCons conservation score (excl. human) |
| GerpN | Neutral evolution score defined by GERP++ |
| GerpS | Rejected Substitution' score defined by GERP++ |
| bStatistic | Background selection score |
| mutIndex | Mutability index from Michaelson, J.J. et al. Cell 2012 |
| dnaHelT | Predicted local DNA structure effect on dnaHelT |
| dnaMGW | Predicted local DNA structure effect on dnaMGW |
| dnaProT | Predicted local DNA structure effect on dnaProT |
| dnaRoll | Predicted local DNA structure effect on dnaRoll |
| EncH3K27Ac | Maximum ENCODE H3K27 acetylation level |
| EncH3K4Me1 | Maximum ENCODE H3K4 methylation level |
| EncH3K4Me3 | Maximum ENCODE H3K4 trimethylation level |
| EncNucleo | Maximum of ENCODE Nucleosome position track score |
| minDistTSS | Distance to closest Transcribed Sequence Start (TSS) |
| minDistTSE | Distance to closest Transcribed Sequence End (TSE) |
| RawScore | Raw score from CADD model |
| PHRED | CADD PHRED Score |

